# Common evolutionary origins of the bacterial glycyl tRNA synthetase and alanyl tRNA synthetase

**DOI:** 10.1101/2023.05.29.542759

**Authors:** Claudia Alvarez-Carreño, Marcelino Arciniega, Lluis Ribas de Pouplana, Anton S. Petrov, Adriana Hernández-González, Marco Igor Valencia-Sánchez, Loren Dean Williams, Alfredo Torres-Larios

## Abstract

Aminoacyl-tRNA synthetases (aaRSs) establish the genetic code. Each aaRS covalently links a given canonical amino acid to a cognate set of tRNA isoacceptors. Glycyl tRNA aminoacylation is unusual in that it is catalyzed by different aaRSs in different lineages of the Tree of Life. We have investigated the phylogenetic distribution and evolutionary history of bacterial glycyl tRNA synthetase (bacGlyRS). This enzyme is found in early diverging bacterial phyla such as Firmicutes, Acidobacteria, and Proteobacteria, but not in archaea or eukarya. We observe relationships between each of six domains of bacGlyRS and six domains of four different RNA-modifying proteins. Component domains of bacGlyRS show common ancestry with i) the catalytic domain of class II tRNA synthetases; ii) the HD domain of the bacterial RNase Y; iii) the body and tail domains of the archaeal CCA-adding enzyme; iv) the anti-codon binding domain of the arginyl tRNA synthetase; and v) a previously unrecognized domain that we call ATL (Ancient tRNA latch). The ATL domain is found only in bacGlyRS and in the universal alanyl tRNA synthetase (uniAlaRS). Further, the catalytic domain of bacGlyRS is more closely related to the catalytic domain of uniAlaRS than to any other aminoacyl tRNA synthetase. The combined data suggest that the ATL and catalytic domains of these two enzymes are ancestral to bacGlyRS and uniAlaRS, which emerged from common protein ancestors by bricolage, stepwise accumulation of protein domains, before the last universal common ancestor of life.

## Introduction

Translation of mRNAs into coded polypeptides is universal to life (1). mRNA decoding during translation depends on the ribosome and a group of enzymes called aminoacyl tRNA synthetases (aaRSs) (2–5). aaRSs determine the genetic code by recognizing specific tRNAs and linking them to their cognate amino acids. aaRSs fall into two classes with independent origins in the deep evolutionary past (6; 7; 4). The catalytic domain of Class I aaRSs is a Rossmann fold while the catalytic domain of Class II aaRSs is an α/β three-layered sandwich. The early evolution of aaRSs has been linked to the origins of the genetic code and to the emergence of the first proteins (8–12).

Throughout the phylogenetic tree, a given tRNA is typically aminoacylated by a single type of aaRS. However, there are exceptions to this rule. There are two distinct lysyl-tRNA synthetases (LysRSs) (13). A class II LysRS is more frequent in bacteria, while a class I LysRS is found predominantly in archaea (14).

There are also two distinct GlyRSs (15–17). The GlyRS found in archaea and eukarya, and in some bacteria, is called here arcGlyRS. The GlyRS found in most in bacteria is called here bacGlyRS. Except for their catalytic domains, bacGlyRS and arcGlyRS are globally different (18) and use different strategies to recognize tRNA^Gly^. arcGlyRS is relatively small, with a catalytic domain and a tRNA recognition domain. bacGlyRS is larger and more complex. Here we ask why there are two GlyRSs. How did they originate? Which GlyRS appeared first? Why is the distribution of the two GlyRSs over the phylogenetic tree patchy?

We and others (19; 17) have hypothesized that the component domains of bacGlyRS were serially exapted through bricolage. The bricolage hypothesis suggests that complex proteins can arise and evolve through stepwise exaptation and adaption of pre-existing domains or modules, which are combined to produce new functions (20). In this hypothesis, domains can be thought of as evolutionary building blocks that provide information on ancestry.

If the bricolage hypothesis is correct, homologies of individual domains can be used to help reconstruct the complex histories of multidomain proteins such as bacGlyRS and can help us understand ancestral relationships. Accurate histories require accurate domain delineations. A domain is an quasi-independent and stable three-dimensional structure (21–23) and is a common unit of evolution (24; 25). Domains are encoded by genes. Duplicated genes, called paralogs (26), can combine to form a variety of distinct multi-domain proteins.

bacGlyRS contains α and β subunits. The combined data are interpreted here to indicate that the β subunit of bacGlyRS holds six component domains rather than five domains as proposed previously (27). The domain of bacGlyRS that escaped detection in previous work is informative about bacGlyRS ancestry and shows homology to a domain found in the universal alanyl tRNA synthetase (uniAlaRS), a class II aaRS. Sequence and structural comparisons suggest that bacGlyRS is more closely related to uniAlaRS than to arcGlyRS (28; 29; 19).

Our results are consistent with a model in which bacGlyRS evolved by bricolage. We show that six component domains of bacGlyRS are homologous to other proteins. These other proteins are universal, bacteria-specific or archaea-specific. bacGlyRS shows common ancestry with six domains of four proteins. Each of these proteins interacts with an RNA although none interact with tRNA^Gly^. Component domains of bacGlyRS show sequence and structural similarities with 1) the catalytic and ATL domains (see below) of uniAlaRS; 2) the anti-codon binding domain (ABD) of the universal arginyl-tRNA synthetase (uniArgRS); 3) the body and tail domains of the archaeal CCA-adding enzyme; and 4) the HD domain of the bacterial RNase Y (in agreement with previous observations of structural similarity (27)). This conservation of RNA as substrate throughout a series of exaptation steps suggests an accessible mechanism by which RNA-associated proteins are repurposed for new RNA-associated functions.

## Results

### bacGlyRS has a complex evolutionary history

The phylogenetic distribution of bacGlyRS appears irregular and discontinuous (‘patchy’) over the extant twigs of the bacterial Tree of Life. We examined a broad range of archaeal and bacterial proteomes, some of which are only partial (Figure 1). Bacterial species have either bacGlyRS or arcGlyRS. Bacteria containing both bacGlyRS and arcGlyRS are extremely rare. Of the 542 sampled proteomes, arcGlyRS (GenBank: PSL06957.1) and bacGlyRS (GenBank: PSL02860.1) are found together only once, in the bacterium *Haloactinopolyspora alba* (NCBI RefSeq assembly: GCA_003014555.1) isolated from sediment of the Dead Sea. Many bacterial phyla have an irregular distribution of the GlyRSs with multiple branches containing either bacGlyRS or arcGlyRS. Firmicutes, Actinobacteria, Chloroflexi, Proteobacteria and DST, PVC, and FCB groups each contain species with bacGlyRS and species with arcGlyRS. Candidate Phyla Radiation (CPR) uniformly contains arcGlyRS; Cyanobacteria uniformly contains bacGlyRS.

**Figure 1.**
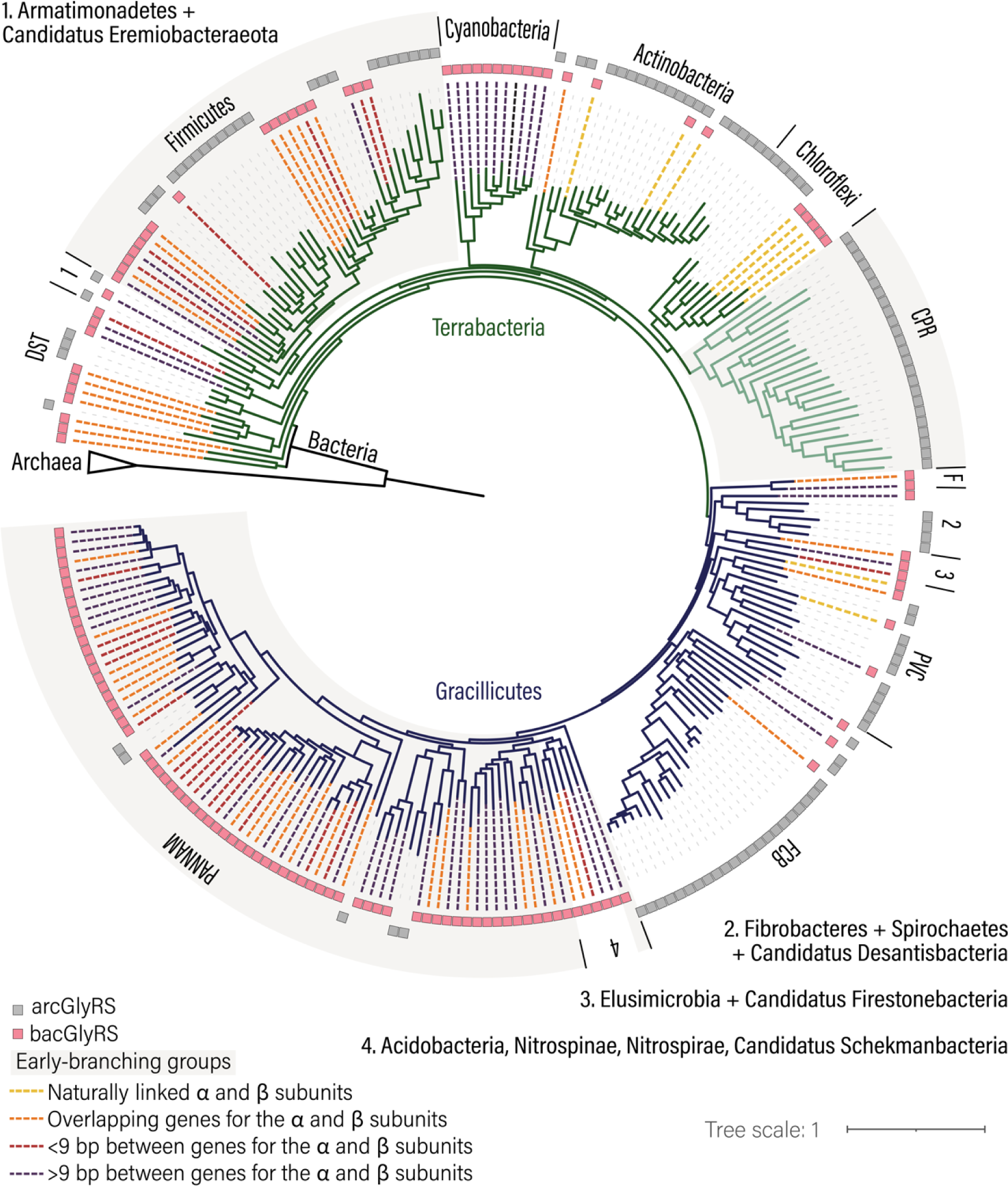
Phylogenetic distribution of the two versions of Glycyl tRNA ligase in Bacteria. The presence of bacGlyRS and arcGlyRS is mapped into a phylogeny of bacterial species. Yellow dotted lines: bacGlyRS with naturally linked α and β subunits form homodimers instead of heterotetramers. Pink dotted lines: species in which the genes for the α and β subunits overlap. Red dotted lines: species in which the genes for the α and β subunits are separated by nine base pairs or less. Purple dotted lines: species in which the genes for the α and β subunits are separated by more than nine base pairs. The topology of the tree was obtained from (32). DST: Deinococcus-Thermus, Synergistetes, Thermotogae, Caldiserica, Coprothermobacterota; CPR: Candidatus Phyla Radiation; PVC: Planctomycetes, Verrucomicrobia, Chlamydiae, Kiritimatiellaeota, Lentisphaerae, Candidatus Desantisbacteria, Candidatus Omnitrophica; FCB: Fibrobacteres, Chlorobi, Bacteroidetes, Gemmatimonadetes, Candidatus Cloacimonetes, Candidatus Fermentibacteria, Candidatus Glassbacteria; PANNAM: Proteobacteria, Aquificae, Nitrospinae, Nitrospirae, Acidobacteria, Chrysiogenetes, Deferribacteres, Schekmanbacteria and Thermodesulfobacteria. F. Fusobacteria. bp: base pairs.

### bacGlyRS sequences are present in early diverging bacteria

The phylogenetic distribution of bacGlyRS suggests that it was present at the Last Bacterial Common Ancestor (LBCA). bacGlyRS is found in early diverging bacterial phyla (30) such as Firmicutes, Acidobacteria, and Proteobacteria (Figure 1). bacGlyRS is not found in the Candidate Phyla Radiation (CPR) phylum. The presence of arcGlyRS in CPR proteomes may be due to horizontal gene transfer rather than vertical inheritance. The CPR phylum contains putative symbiotic and parasitic bacteria with reduced genomes, large gene losses and extensive lateral gene transfer (31).

### Domains in bacGlyRS are related to RNA-modifying enzymes

Each domain of bacGlyRS has an independent evolutionary history. Six domains in bacGlyRS display strong sequence similarity to domains of other proteins (Figure 2). We determined proteins that are the closest sequence matches to each of the six domains of bacGlyRS (Figure 2E). The catalytic domain shows sequence similarity to the catalytic domain of uniAlaRS (e-value 1.1×10^−07^). The body and tail domains show sequence similarity to domains of the archaeal CCA-adding enzyme (e-value 1.4×10^−08^). The HD domain shows sequence similarity to a domain of ribonuclease Y (RNase Y) (e-value 5.7×10^−08^). The ABD shows sequence similarity to a domain of uniArgRS (e-value 2.9×10^−17^).

**Figure 2.**
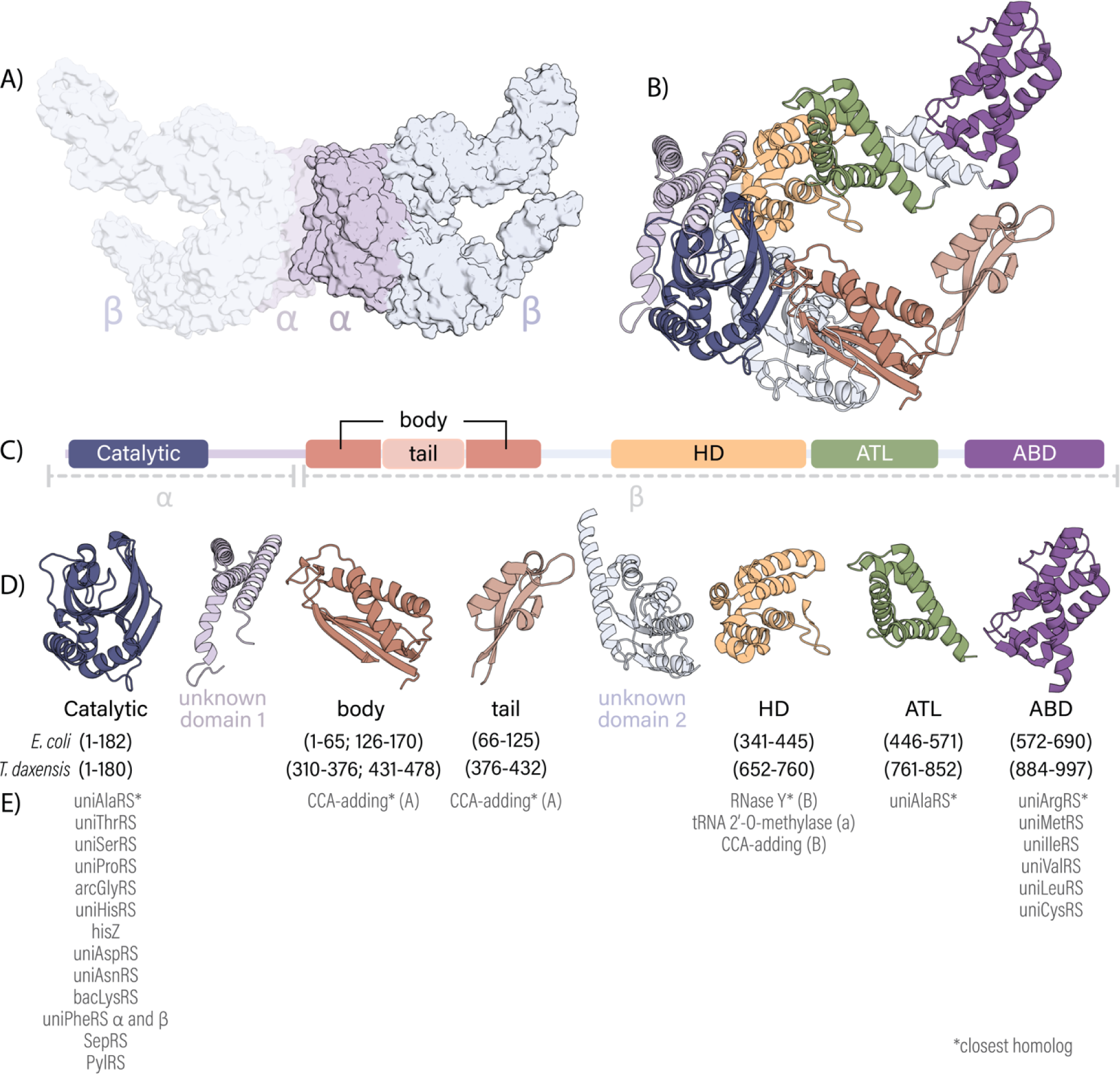
Domain organization of bacGlyRS. Structure of bacGlyRS of *Thermoanaerotrix daxensis* with naturally linked α and β-subunits (PDB: 7LU4). (A) Surface representation of bacGlyRS indicating the region that corresponds to the α-(light purple) and β-(light blue) subunits in other bacteria. (B) A ribbon representation of bacGlyRS shows structural units in different colors: dark blue, catalytic domain; light purple, unknown domain 1; dark orange, body domain; light orange, tail domain; light blue, unknown domain 2; yellow, HD domain; green, ATL domain; and dark purple, ABD domain. (C) Domains and structural units of bacGlyRS. (D) Proteins containing domains with high-scoring sequence similarity by bidirectional BLAST.

We identified a previously unrecognized domain in bacGlyRS that we call Ancient tRNA latch (ATL) domain. Our determination that ATL is a distinct domain follows previous identification of this domain in uniAlaRS (33). The ATL domains of uniAlaRS and bacGlyRS are similar to each other in sequence and structure (Figure 3A). If ATL is a distinct domain in uniAlaRS, then it is also a domain in bacGlyRS. Previously, the ATL domain was described incorrectly as a dependent element of the HD domain (27; 17; 34); we infer here from sequence and structural considerations that the ATL is independent of other elements of bacGlyRS. The ATL domain appears to be unique to bacGlyRS and uniAlaRS, no other homologs of the ATL domain are identified in a comprehensive protein sequence database (UniRef90). The ATL domain of uniAlaRS contains a C-terminal α-helical decoration that is absent from the ATL domain of bacGlyRS.

**Figure 3.**
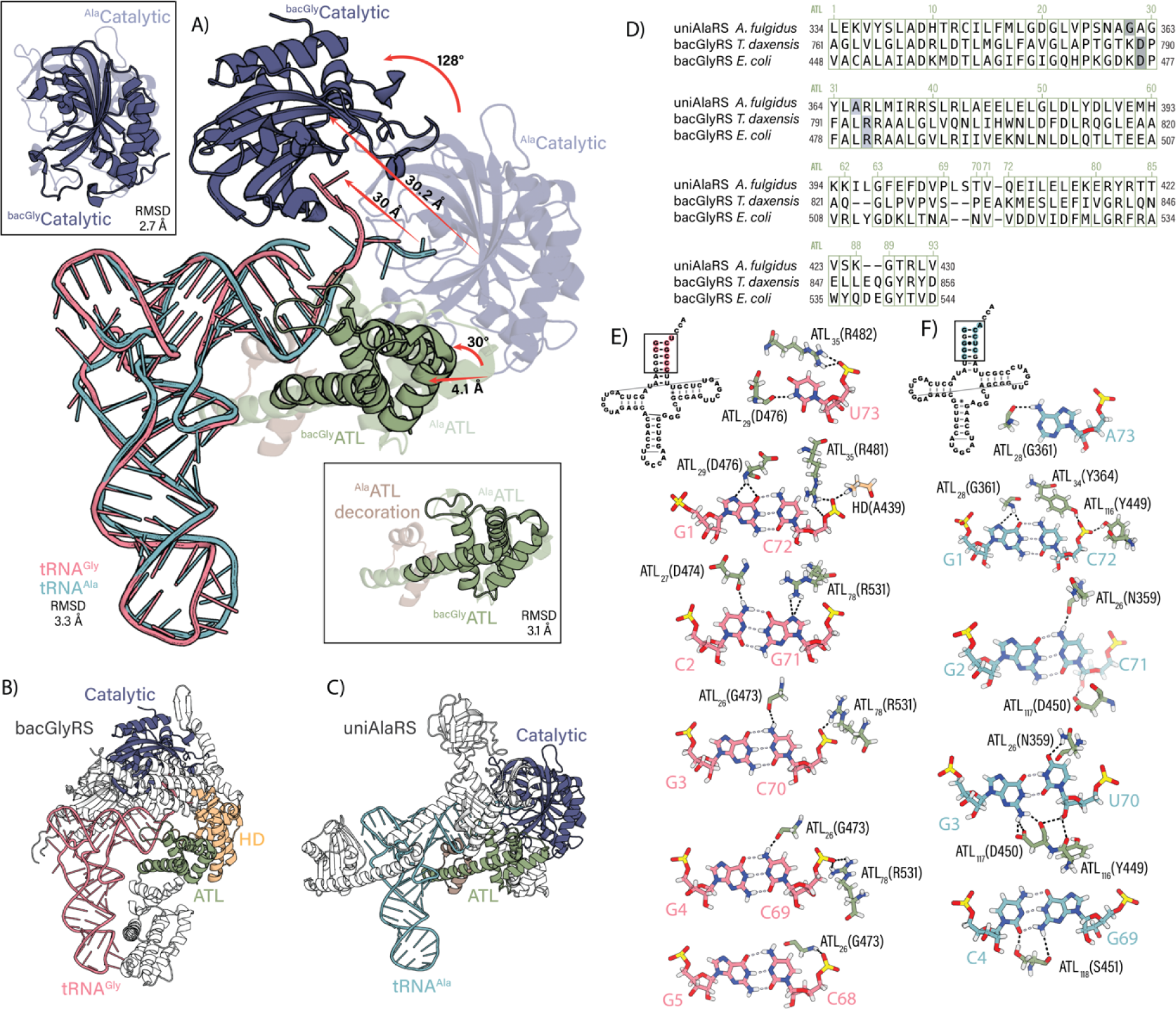
Structures of bacGlyRS and uniAlaRS showing regions of homology. (A) Comparison of the relative position of the catalytic and ATL domains of bacGlyRS (PDB: 7YSE) and uniAlaRS (PDB: 3WQY) with respect to their cognate tRNA molecules. Insets: Structure superimposition of the catalytic and ATL domains of bacGlyRS and uniAlaRS. (B) bacGlyRS of *Escherichia coli* in complex with tRNA. (C) uniAlaRS of *Archaeoglobus fulgidus* in complex with tRNA. (D) Sequence alignment and indexing of the ATL domains of uniAlaRS and bacGlyRS. (E) Hydrogen bonds between the ATL domain of bacGlyRS and its cognate tRNA. (F) Hydrogen bonds between the ATL domain of uniAlaRS and its cognate tRNA.

In addition, we observe that the function of the ATL domain has been mischaracterized. The ATL domain in uniAlaRS is called the “putative anticodon-binding domain” in some classification databases (SCOPe ID a.203.1.1, (35), and ECOD ID 613.1.1.1 (36)). However, neither the ATL domain of uniAlaRS nor the ATL domain of bacGlyRS contacts the anti-codon stem-loop of tRNA (37; 38). Both interact extensively with the amino acid acceptor stem. This mischaracterization of the ATL domain structure and function has obscured its evolutionary origins and molecular function.

### bacGlyRS and uniAlaRS homology is observed in key functional domains

The primary functions of aaRSs are to recognize tRNAs and amino acids and to covalently join them together. The catalytic and ATL domains, which carry out these essential functions are homologous between bacGlyRS and uniAlaRS.

The ATL domains of uniAlaRS and bacGlyRS recognize tRNAs by variations of a common mechanism. tRNAs are recognized by uniAlaRS and bacGlyRS by interactions of ATL with the tRNA acceptor stem (27; 17; 34). The tRNA acceptor stem is the first seven base pairs of tRNA, the unpaired nucleotide at position 73, and the 3’-CCA tail. In species containing bacGlyRS, the acceptor stems of tRNA^Gly^ and tRNA^Ala^ differ in three pairs and in position 73. These three base pairs and position 73 are signatures (39), meaning that they are highly conserved within each group, tRNA^Gly^ or tRNA^Ala^, but differ between the two groups. The specific signatures of tRNA^Gly^ are base pairs C2:G71, G3:C70, G4:C69, and U73. The signatures of tRNA^Ala^ are G2:C71, G3:U70, C4:G69, and A73. In uniAlaRS, the wobble base pair G3:U70 is the main identity determinant for tRNA^Ala^ aminoacylation by uniAlaRS (40; 33; 37; 41; 42; 34). The ATL domain interacts with the base of the discriminator position (U73 in tRNA^Gly^, and A73 in tRNA^Ala^).

The ATL domain has adapted to distinguish features of the amino acid acceptor stems of tRNA^Gly^ or tRNA^Ala^ (Figure 3). We have identified four distinct ATL adaptations, allowing a common fold to recognize different tRNAs. To discriminate between the two different tRNAs, the ATL domain (i) slightly shifts global position relative to the tRNA, (ii) shifts local position of the secondary structural elements by conformational change, (iii) changes sequence and (iv) acquires α-helical decorations.

The ATL domains of bacGlyRS and uniAlaRS exhibit slightly different positions relative to the tRNA. A distance of 4.1 Å between the centers of mass of ^bacGly^ATL and ^Ala^ATL after direct superimpositions of the tRNA^Gly^ in complex with bacGlyRS (PDB 7YSE) and the tRNA^Ala^ in complex with uniAlaRS (PDB 3WQY), indicates the changes in the global pose of the ATL domains (Figure 3A).

The interactions between ATL and tRNA are locally different in ^bacGly^ATL and ^Ala^ATL. After direct superimposition of the ^bacGly^ATL and ^Ala^ATL protein domains, a 3.1 Å root mean square deviation (RMSD) between the superimposed backbone atoms quantitates local changes in conformation (Figure 3A). To describe local positional shifts, we have reindexed the ATL residues so that a common index (subscripted here) indicates the same column in the sequence alignment (Figure 3D). ^bacGly^ATL_10_ indicates the amino acid at the 10^th^ alignment position of ATL domain of bacGlyRS while ^Ala^ATL indicates the amino acid of the ATL domain of uniAlaRS at the same alignment position. Differences in local positioning of amino acids within ATL are seen by differences in ATL indexes of a common interaction. For example, local positional shifts are seen in the interaction between ^bacGly^ATL and ^Ala^ATL with the G1 base and the discriminator positions of the tRNAs (U73 in tRNA^Gly^ and A73 in tRNA^Ala^) and in the interaction between ^bacGly^ATL and ^Ala^ATL with C72 and the discriminator. Base pair G1:C72 is conserved in both tRNA^Gly^ and tRNA^Ala^. By contrast, there is no shift at index 26. Both ^bacGly^ATL and ^Ala^ATL interact with position 70, which is a C in tRNA^Gly^ and a U in tRNA^Ala^.

The ^bacGly^ATL interaction with tRNA appears intrinsically less stringent than that of ^Ala^ATL. The ^bacGly^ATL domain contacts nucleotides of only one strand in the region of C68, C69 and C70. It does not contact the opposing strand (G3, G4 or G5, Figure 3E). By contrast, ^Ala^ATL targets both strands of tRNA^Ala^, interacting with both bases of base-pairs G3:U70 and C4:G69 in part by using the ^Ala^ATL-specific C-terminal decoration (Figure 3F).

The catalytic domain undergoes a radical change in global position between bacGlyRS and uniAlaRS. In uniAlaRS, the catalytic and ATL domains are directly adjacent in sequence and structure. By contrast, in bacGlyRS, the HD and body-tail domains are inserted between the catalytic and ATL domains. The HD domain shifts and rotates the catalytic domain of bacGlyRS by 130° compared to the catalytic domain of uniAlaRS (Figure 3B). To accommodate the change in the position of the catalytic domain of bacGlyRS, the tRNA distorts. The conformation of the four unpaired nucleotides of the acceptor arm of tRNA^Gly^ bends (Figure 3A), such that the position of the terminal adenosine is shifted by ∼ 26 Å in the bacGlyRS complex compared to in the uniAlaRS complex.

The conformation of the CCA tail and the position of the terminal adenosine appear to be unique in bacGlyRS compared to other aaRSs. The CCA tail of tRNA^Gly^ bends away from the anti-codon in the complex with bacGlyRS (34), shifting the terminal A toward the minor groove of the tRNA. All other class II aaRSs bend the CCA tail in the opposite direction, towards the anticodon (Figure S1) or, in the case of the uniAlaRS-tRNA complex (Figures 3A and 3C), maintain the relaxed linear structure (43). The position of the terminal adenosine is shifted by ∼ 30 Å in the tRNA complex of bacGlyRS compared to in the PheRS complex (Figure S1).

### The catalytic and ATL domains of bacGlyRS resemble archaeal uniAlaRS homologs

We formulated a tentative model in which bacGlyRS originated in bacteria via duplication of a two-domain uniAlaRS module. One domain is ATL and the other is the catalytic domain. This tentative model predicts that bacGlyRS component domains would be most similar to bacterial uniAlaRS homologs. Therefore, we investigated the similarity of bacGlyRS domains to archaeal and bacterial uniAlaRS sequences. As shown here this model is not supported by the data.

The sequences of the catalytic and ATL domains in bacterial uniAlaRS and archaeal uniAlaRS are distinctive. A composite multiple-sequence alignment (MSA) containing both bacterial and archaeal uniAlaRS sequences reveals lineage-specific signature positions and insertions (Figure S2). Signature positions correspond to columns in a composite MSA that are conserved within groups but are different between groups (39). Thus, differences between bacterial and archaeal uniAlaRS sequences reflect deep divergence between bacteria and archaea and are not due to a simple bias in amino acid usage. This type of pattern in a composite MSA is called a block structure and is observed in many universal aaRS and other proteins of the translation system (3; 44).

The catalytic and ATL domains of bacGlyRS resemble archaeal uniAlaRS homologs more closely than bacterial uniAlaRS homologs (Figures 4A and B). We compared the sequences of the catalytic and ATL domains of bacGlyRS with those of uniAlaRS using the program CLANS (45; 46) and their profiles using the program HH-align (47). Profiles are calculated from multiple-sequence alignments and reflect position-specific sequence variability. A cluster map of sequences from the catalytic domains of bacGlyRS and uniAlaRS at a P-value lower than 10^−12^ shows connections between bacGlyRSs and archaeal uniAlaRSs. At this P-value threshold, the cluster map lacks connections between bacGlyRSs and bacterial uniAlaRSs (Figure 4A). Similarly, a cluster map of sequences from the ATL domains of bacGlyRS and uniAlaRS at a P-value threshold of 10^−11^ shows connections between bacGlyRSs and archaeal uniAlaRSs. This cluster map lacks connections between bacGlyRSs and bacterial uniAlaRSs (Figure 4B). Thus, the catalytic and ATL domains of bacGlyRS are more similar to archaeal homologs than to bacterial homologs. Therefore, our tentative model in which bacGlyRS originated by duplication of a bacterial uniAlaRS module composed of catalytic and ATL domains (Figure 5D) is not supported by the data (Figure 4).

**Figure 4.**
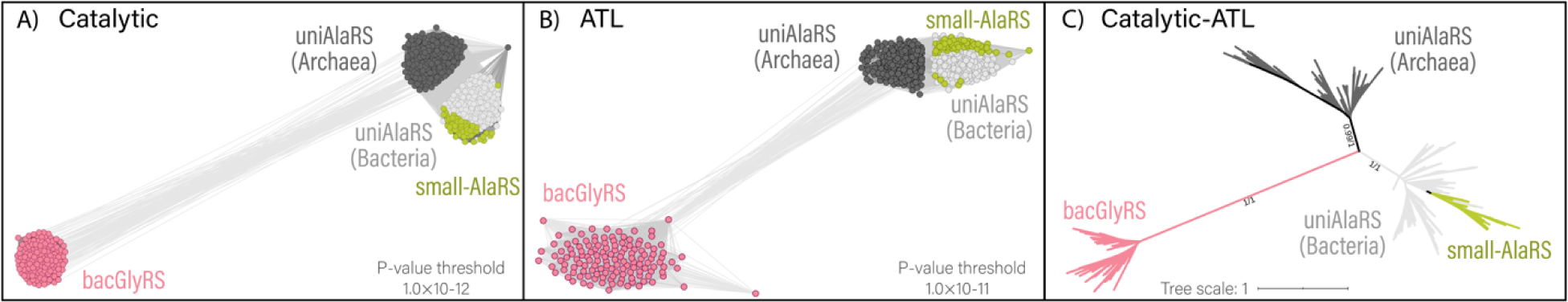
Sequence analysis of bacGlyRS protein domain homologs. Cluster map based on pairwise sequence similarities. Nodes represent sequences; connections between nodes indicate matches between pairs of sequences with a *P*-value lower than the threshold. (A) Pairwise sequence similarity between the catalytic domains of bacGlyRS and uniAlaRS. (B) Pairwise sequence similarity between the ATL domains of bacGlyRS and uniAlaRS. (C) Maximum-likelihood (LG+R4) unrooted tree of catalytic and ATL domains including bacGlyRS and uniAlaRS sequences. small-AlaRS are sequences from an anomalous subgroup of uniAaRS found in some archaea and bacteria.

**Figure 5.**
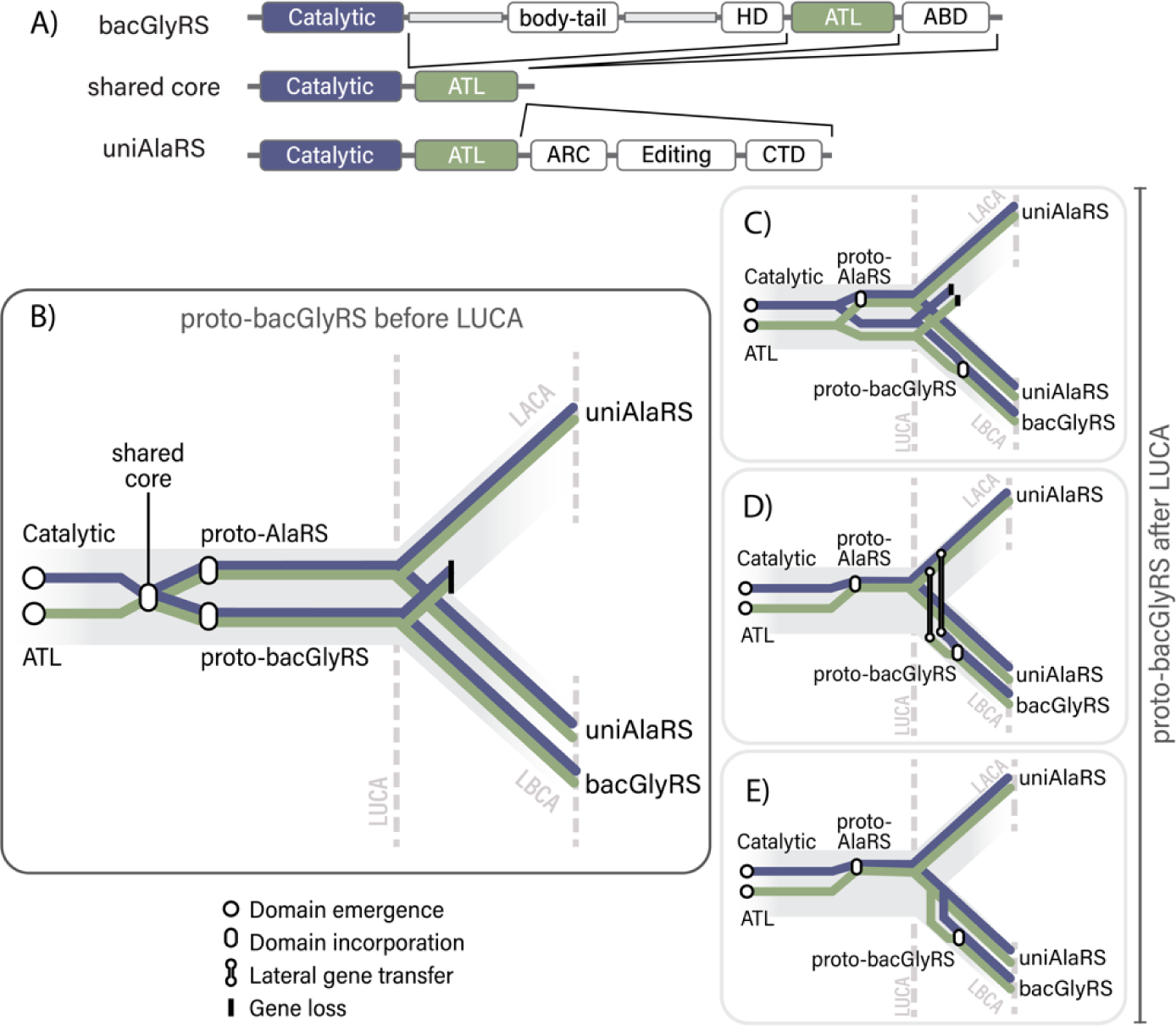
Emergence of bacGlyRS through molecular bricolage. (A) Comparative diagram of the multidomain arrangements of bacGlyRS and uniAlaRS. (B) Emergence of the common ancestor of bacGlyRS and uniAlaRS before the divergence of LUCA. Duplication of the common ancestor gives rise to the proto-uniAlaRS and proto-bacGlyRS. The ancestor of bacGlyRS is lost in archaea. (C) Emergence and duplication of the catalytic and ATL domains at LUCA. Before the divergence between archaea and bacteria, one copy of the catalytic domain and one copy of the ATL domain form the uniAlaRS core, the paralogs of the catalytic and ATL domains continue as single-domain proteins. Secondary loss of the copy of the catalytic and ATL domains in archaea. Formation of the bacGlyRS core in bacteria. (D and E) Evolution of the core of uniAlaRS before the divergence of LUCA. (D) The catalytic and ATL domains are horizontally transferred from the archaeal to the bacterial lineage and formation of the bacGlyRS core. (E) Duplication of the catalytic and ATL domains from uniAlaRS in bacteria and formation of the bacGlyRS core.

### The catalytic and ATL domains represent an evolutionary and functional core

Canonical uniAlaRS has five component domains; (i) the catalytic domain, which aminoacylates the tRNA, (ii) the ATL domain, which recognizes tRNA^Ala^, (iii) a cradle loop barrel domain, (iv) an editing domain and (v) a C-terminal domain (C-Ala) that binds to the tRNA elbow formed by the D- and T-loops (48; 33) (Figures 3C and S1). We identified a subgroup of bacterial and archaeal uniAlaRS sequences that have only three of these domains: catalytic, ATL, and editing (Figures 4 and S1). uniAlaRSs in this subgroup lack the cradle loop barrel domain and the C-Ala domain of canonical uniAlaRSs (Figure S2). We call this anomalous subgroup small-AlaRS. We identified small-AlaRS in the proteomes of the CPR bacterium *Candidatus Thermoplasmatota*, and in DPANN, and Asgard groups. small-AlaRS proteins have bacteria-specific insertions, suggesting that sequences were transferred to some archaea via horizontal gene transfer. The maximum likelihood tree of catalytic and ATL domain sequences supports a horizontal transfer hypothesis of small-AlaRS to some archaea (Figure 4C and S3). Small-AlaRS sequences and recently discovered viral mini-AlaRS sequences (49), which lack the cradle loop barrel, the editing and the C-Ala domains of canonical cellular uniAlaRSs, support the hypothesis that a core composed of the catalytic and ATL domains can be functional.

The combined data support a model in which bacGlyRS and uniAlaRS emerged before LUCA from common catalytic and ATL ancestors (Figure 5A and B). These catalytic and ATL ancestors were single-domains and were independent of each other. Other possibilities put the emergence of the bacGlyRS core after the divergence of LUCA. This late emergence of bacGlyRS appears less parsimonious because it suggests that some bacteria lost the ancestral GlyRS and replaced it with an RS protein that has no homologs in other bacteria. Possible scenarios are 1) formation of the proto-bacGlyRS in the bacterial lineage independently from the uniAlaRS catalytic-ATL core (Figure 5C); 2) formation of the proto-bacGlyRS by partial or complete duplication of the uniAlaRS gene in the archaeal lineage, followed by a lateral gene transfer of the proto-bacGlyRS from archaea to the bacterial lineage; and 3) formation of the proto-bacGlyRS by partial or complete duplication of the uniAlaRS gene in the bacterial lineage (Figure 5D). This last scenario is not supported by the data (Figure 4).

## Discussion

bacGlyRS has a deep and convoluted evolutionary history. We infer exaptation relationships (50) between six bacGlyRS domains and six domains of four different RNA-modifying proteins (Figure 2). Exaptation is the co-opting of an existing molecule, domain, trait or system for new function. In our model, upon or after fusion of the ATL and catalytic domains to form ancestors of both bacGlyRS and uniAlaRS, the catalytic domain was exapted to link glycine and tRNA^Gly^ to form glycyl-tRNA^Gly^. The ATL domain was exapted to recognize the acceptor-stem of the tRNA^Gly^ (17). The prior function of the catalytic domain may have been non-specific linkage of amino acids to RNAs. The prior function of ALT is unknown. After these processes, insertion of the HD, body and tail domains impacted the relative positions of the ALT and catalytic domains of bacGlyRS and the interaction of the catalytic domain with the CCA tail of tRNA^Gly^ (see below). The HD domain was exapted to interact with the discriminator position of tRNA^Gly^ (17). The body and tail domains of bacGlyRS were exapted to interact with the arm and elbow of tRNA^Gly^ (34). In the archaeal CCA-adding enzyme, the body and tail domains also interact with the acceptor stem of the tRNA (51; 52). The ABD domain, which is homologous to a domain in the class I uniArgRS, and shares ancestry with it, was exapted to interact with the anti-codon loop of tRNA^Gly^ (34).

Here we provide evidence that the ATL domain is a distinct, independent domain of bacGlyRS rather than an integral component of the HD domain as previously proposed. This distinction is important for understanding the deep evolutionary history of aaRSs. Homology between the catalytic and ATL domains of bacGlyRS and the corresponding domains of uniAlaRS suggests common ancestry of the core domains and functions of these enzymes. It is likely that an ancestral class II catalytic domain and an ATL domain joined, and the resulting two-domain aaRS gave rise to two separate enzymes, one for glycine and another for alanine. This simple model is consistent with sequence and structural similarities between the core domains of bacGlyRS and uniAlaRS.

The conservation of the pose of the ATL domain in uniAlaRS and bacGlyRS, but not the pose of the catalytic domain, is consistent with a model in which the ATL domain is ancestral to both uniAlaRS and bacGlyRS (Fig. 5B). It appears that the ATL domain played a role in primitive RNA biology before it was incorporated into these two synthetases. We have identified four adaptations of ATL that allowed a common fold in a common pose to recognize different tRNAs. The ATL domain (i) slightly shifts global position relative to the acceptor stem of the tRNA, (ii) shifts local positions of amino acid backbone and sidechains by conformation change, (iii) changes sequence, and (iv) acquires a C-terminal decoration in the uniAlaRS that is absent from the bacGlyRS.

bacGlyRS and uniAlaRS use different strategies to achieve final recognition of their cognate tRNAs (Figure 3). ATL domains of both enzymes have similar interactions with the discriminator base and the G1:C72 base pair which is conserved in tRNA^Gly^ and tRNA^Ala^. uniAlaRS uses the ATL domain to interact intensively with the acceptor stem, employing the C-terminal decoration to assist in targeting both strands of the acceptor step. The ATL domain in bacGlyRS lacks C-terminal decoration and interacts less intensively with the acceptor stem. In the bacGlyRS complex, the ATL domain only contacts nucleotides of one strand of the acceptor stem. To compensate, bacGlyRS interacts with other regions of the tRNA. bacGlyRS but not uniAlaRS exapted additional recognition domains through a process of bricolage.

The conformation of the CCA tail and the position of the terminal adenosine appear to be unique in bacGlyRS compared to other aaRSs. In uniAlaRS, the catalytic and ATL domains are directly adjacent to each other in sequence and structure. tRNA^Ala^ interacts with the CCA tail of uniAlaRS in a relaxed state, which is reasonably similar to the structure in the free tRNA (53). In bacGlyRS, insertion of the HD domain caused a translation and rotation of the catalytic domain relative to the ATL domain (Figure 3B). To accommodate this translation/rotation, the acceptor-arm of tRNA^Gly^ in the bacGlyRS complex bends acutely in the direction of the minor groove (Figure 3C). This conformation of the tRNA appears to be unique to bacGlyRS.

The sequences of bacGlyRS and archaeal uniAlaRS share distinctive features. The catalytic and ATL domains of bacGlyRS more closely resemble uniAlaRS archaeal homologs than bacterial homologs (Figures 4A and B). Thus, a model in which bacGlyRS originated within the bacterial lineage by duplication of a bacterial catalytic-ATL uniAlaRS module is not supported by the data (Figure 5D). The combined results suggest that bacGlyRS and uniAlaRS share a common ancestor composed of the catalytic and ATL domains (Figure 5A). The catalytic and ATL domains represent an evolutionary and functional core. The domains in the shared common core of bacGlyRS and uniAlaRS interact with the most ancient region of the tRNA (54; 55).

Here we propose a simple model, in which insertion of the body-tail and HD domains (or possibly just the HD domain) into a gene closely related to the proto-AlaRS formed a novel enzyme, a proto-bacGlyRS (Figure 5A). In this model, the interaction between ATL domain and the tRNA remained relatively unchanged as the catalytic domain was displaced and the CCA tail bends towards the minor groove. The proto-bacGlyRS enzyme underwent adaptations that conferred a new amino acid specificity (Gly) and a new tRNA identity (tRNA^Gly^). It is possible that the new tRNA was better suited to the CCA tail orientation imposed by the HD insertion.

bacGlyRS is one of the largest aaRSs and is usually composed of two subunits, with a catalytic domain in the α subunit and several recognition domains in the β subunit. In most bacteria (Figure 1) bacGlyRS is a heterotetramer (4). In some bacteria, including *Thermoanaerotrix daxensis*, the α and β subunits are linked in a single polypeptide chain (Figure 1); bacGlyRS with linked α and β subunits form homodimers (Figure 2A). The form of oligomerization, the presence of α-β linked bacGlyRSs and the encoding of α and β bacGlyRS subunits as a single (56; 57), and sometimes overlapping (Figure 1, Table S1), open reading frame, could indicate that the ancestor of all bacGlyRSs was a single α-β protein that formed homodimers. Fragmentation of bacGlyRS may have occurred later (57) and persists in most species due to evolutionary pressures (27). Secondary fragmentation of uniAlaRS has also been reported (58).

The distribution of bacGlyRS in a wide range of deeply rooted bacterial species suggests origins before LBCA. The presence of bacGlyRS in Terrabacteria and Gracilicutes, the two major bacterial groups (Figure 1) also supports this early origin. However, in the bacterial tree the distribution of bacGlyRS and arcGlyRS appears patchy and irregular. The dominance of arcGlyRS in CPR proteomes suggests that symbiotic bacteria with reduced genomes prefer a smaller enzyme. Currently, we do not have a model that fully explains the discontinuous distribution over evolutionary history.

Our observations here reflect deep evolutionary history. Glycine and alanine appear to be ancient amino acids, both prebiotically available and biologically important. They are among the most abundant amino acids produced in spark discharge experiments (59) and are found in chondrite asteroids (60). We propose that bacGlyRS and uniAlaRS emerged from a shared core before LUCA (Figure 5A and B), from catalytic and ATL domains that had separate and distinct origins. These prebiotic amino acids share the first and third codon positions, and the proteins that aminoacylate tRNA^Gly^ and tRNA^Ala^ share sequence and structure relationships. It appears we are seeing evidence of an ancestral protein with non-specific catalytic activity that may have been able to link either glycine or alanine to an RNA, presumably an ancestor of tRNA. The deep relationship between alanine and glycine is seen in their shared chemistry and biology.

## Methods

### Sequence similarity search

A multiple sequence alignment (MSAs) of bacGlyRS sequences was calculated for a representative set of archaea and bacteria. The MSA was trimmed to the domain boundaries as defined by the structure analysis. The MSAs of the domains were converted to profile Hidden Markov Models using the HHsuite version 3.3.0 (31). A profile-profile similarity search on the Evolutionary Classification of Domains (ECOD) database was performed with HHsearch using the profiles of bacGlyRS domains as queries. Target domains in ECOD yielding HHsearch probabilities greater than 60% over more than 50 aligned columns were retrieved to serve as queries of a second search on a sequence database containing the proteomes of the representative set of bacterial and archaeal species. The second search was performed using jackhmmer from the HMMER suite version 3.3.1 (31), with default parameters (five iterations). The ECOD database only contains domains that are directly annotated on proteins with experimentally determined structures. The second search aims to identify sequence similarity in the UniProt Reference Cluster, UniRef90 (61); this database includes proteins without experimentally determined structures. Sequences passing the threshold values (E-value < 1×10^−3^, query coverage 80%) were retrieved and trimmed to the region of sequence similarity.

We broadly describe the phylogenetic distribution of the best hits according to the number of sampled proteomes containing homologous sequences as follows: universal (presence in > 50% of archaea and >50% of bacteria); archaea-specific (presence in >50% of archaea and <10% of bacteria); bacteria-specific (presence in < 10% of archaea and <50% of bacteria); lineage-specific within archaea (presence in <50% of archaea and <50% of bacteria); lineage-specific within bacteria (presence in <10% of archaea and <50% of bacteria sampled); or of complex distribution (presence in 10-50% of archaea and 10-50% of bacteria).

### Cluster analysis

The query MSAs and trimmed target sequences retrieved from the second sequence similarity search were clustered with CLANS (46) based on pairwise sequence similarities. To find the best sequence match of each query domain, the cut-offs of the clustering were adjusted to find the P-value for which the group of query sequences shows sequence relationships with only one other group of orthologs. The subset of best matches was further clustered to reveal possible differences between archaeal and bacterial sequences. Bacterial and archaeal uniAlaRS sequences displayed differential similarity to bacGlyRS sequences on the cluster analysis. To corroborate the clustering analysis results, additional analyses were performed at the level of profiles.

### Pairwise profile analysis: HHalign

An MSA containing orthologous sequences of uniAlaRS was further divided to include only archaeal sequences or only bacterial sequences. The MSAs of archaeal and bacterial sequences of uniAlaRS and the MSA of bacGlyRS were trimmed to the catalytic and ATL domains. The MSA of the catalytic domain of bacGlyRS was aligned to the MSAs of the catalytic domains of archaeal and bacterial uniAlaRS sequences using HHalign. The HHalign scores of these MSA-MSA comparisons agree with the differential pairwise similarities observed in CLANS between archaeal and bacterial sequences.

### ML tree

The composite MSA of the catalytic domain containing sequences from bacGlyRS and archaeal sequences from uniAlaRS was concatenated to the composite MSA of the ATL domain. The concatenated MSA was trimmed with trimAl (62). A Maximum Likelihood tree was calculated using the trimmed MSA using PhyML (63) and SMS model selection tool (64) on the Montpellier Bioinformatics Platform. The tree was inferred using an LG+R4 model of evolution. Branch support corresponds to a Bayesian-like transformation of aLRT (aBayes) (65).

## Data Availability

Sequence alignments and CLANS input files associated with this manuscript have been deposited in the FigShare repository DOI: 10.6084/m9.figshare.23269007.

## Abbreviations and symbols

aaRS: aminoacyl tRNA synthetase
ABD: anti-codon binding domain
arcGlyRS: glycyl aminoacyl tRNA synthetase of archaea and eukarya, and some bacteria
ATL: Ancient tRNA latch domain
bacGlyRS: bacterial glycyl aminoacyl tRNA synthetase
CPR: Candidate Phyla Radiation
GlyRS: glycyl aminoacyl tRNA synthetase
LysRS: Lysyl aminoacyl tRNA synthetase
mRNA: messenger RNA
MSA: multiple-sequence alignment RNase
Y: ribonuclease Y
tRNA: transfer RNA
uniAlaRS: universal alanyl tRNA synthetase
uniArgRS: universal arginyl-tRNA synthetase

## Acknowledgements

The authors dedicate this work to the memory of Professor ATL who passed away on September 9th, 2021. Professor ATL worked on the structural determination of the bacterial glycyl tRNA synthetase and performed the initial structural analysis. The authors thank Jessica Bowman, Eric Smith and Arturo Becerra for discussions. This work was funded by the National Aeronautics and Space Administration grant 80NSSC18K1139. CAC was sponsored by the National Aeronautics and Space Administration (NASA) through a contract with ORAU.

## Author contributions

Conceptualization, ATL, MIVS and CAC. Methodology, CAC. Formal analysis CAC, MA, AHG, ASP, MIVS and LDW. Writing – Original Draft, CAC and LDW. Writing – Review & Editing, ASP, LRP and MIVS.

## References

1. Bowman JC, Petrov AS, Frenkel-Pinter M, Penev PI, Williams LD (2020) Root of the Tree: The Significance, Evolution, and Origins of the Ribosome. Chem Rev 120:4848–4878.

2. Burbaum JJ, Schimmel P (1991) Structural relationships and the classification of aminoacyl-tRNA synthetases. J Biol Chem 266:16965–16968.

3. Woese CR, Olsen GJ, Ibba M, Söll D (2000) Aminoacyl-tRNA synthetases, the genetic code, and the evolutionary process. Microbiol Mol Biol Rev 64:202–236.

4. Gomez MAR, Ibba M (2020) Aminoacyl-tRNA synthetases. RNA 26:910–936.

5. Giegé R, Eriani G (2023) The tRNA identity landscape for aminoacylation and beyond. Nucleic Acids Res 51:1528–1570.

6. Eriani G, Delarue M, Poch O, Gangloff J, Moras D (1990) Partition of tRNA synthetases into two classes based on mutually exclusive sets of sequence motifs. Nature 347:203–206.

7. Moras D (1992) Structural and functional relationships between aminoacyl-tRNA synthetases. Trends Biochem Sci 17:159–164.

8. Hartman H (1995) Speculations on the evolution of the genetic code IV the evolution of the aminoacyl-tRNA synthetases. Orig Life Evol Biosph 25:265–269.

9. Ribas de Pouplana L, Schimmel P (2001) Aminoacyl-tRNA synthetases: potential markers of genetic code development. Trends Biochem Sci 26:591–596.

10. Fournier GP, Andam CP, Alm EJ, Gogarten JP (2011) Molecular evolution of aminoacyl tRNA synthetase proteins in the early history of life. Orig Life Evol Biosph 41:621–632.

11. Koonin EV, Novozhilov AS (2017) Origin and evolution of the universal genetic code. Annu Rev Genet 51:45–62.

12. Ribas de Pouplana L. The evolution of aminoacyl-tRNA synthetases: From dawn to LUCA. (2020) The Enzymes. Elsevier, pp. 11–37.

13. Terada T, Nureki O, Ishitani R, Ambrogelly A, Ibba M, Söll D, Yokoyama S (2002) Functional convergence of two lysyl-tRNA synthetases with unrelated topologies. Nat Struct Biol 9:257–262.

14. Tumbula D, Vothknecht UC, Kim H-s, Ibba M, Min B, Li T, Pelaschier J, Stathopoulos C, Becker H, Söll D (1999) Archaeal aminoacyl-tRNA synthesis: diversity replaces dogma. Genetics 152:1269–1276.

15. Ostrem DL, Berg P (1974) Glycyl transfer ribonucleic acid synthetase from Escherichia coli: purification, properties, and substrate binding. Biochemistry 13:1338–1348.

16. Tang SN, Huang JF (2005) Evolution of different oligomeric glycyl-tRNA synthetases. FEBS Lett 579:1441–1445.

17. Han L, Luo Z, Ju Y, Chen B, Zou T, Wang J, Xu J, Gu Q, Yang X-L, Schimmel P (2023) The binding mode of orphan glycyl-tRNA synthetase with tRNA supports the synthetase classification and reveals large domain movements. Science Advances 9:eadf1027.

18. Logan D, Mazauric M, Kern D, Moras D (1995) Crystal structure of glycyl-tRNA synthetase from Thermus thermophilus. EMBO J 14:4156–4167.

19. Dimas-Torres J, Rodríguez-Hernández A, Valencia-Sánchez M, Campos-Chávez E, Godínez-López V, Rodríguez-Chamorro D, Grøtli M, Fleming C, Hernández-González A, Arciniega M (2021) Bacterial glycyl tRNA synthetase offers glimpses of ancestral protein topologies.

20. Jacob F (1977) Evolution and tinkering. Science 196:1161–1166.

21. Rossman MG, Liljas A (1974) Letter: Recognition of structural domains in globular proteins. J Mol Biol 85:177–181.

22. Levitt M, Chothia C (1976) Structural patterns in globular proteins. Nature 261:552–558.

23. Porter LL, Rose GD (2012) A thermodynamic definition of protein domains. Proc Natl Acad Sci USA 109:9420–9425.

24. Ponting CP, Schultz J, Copley RR, Andrade MA, Bork P (2000) Evolution of domain families. Advances in protein chemistry 54:185–244.

25. Ohno S. 2013. Evolution by gene duplication, Springer Science & Business Media.

26. Fitch WM (1970) Distinguishing homologous from analogous proteins. Syst Zool 19:99–113.

27. Ju Y, Han L, Chen B, Luo Z, Gu Q, Xu J, Yang XL, Schimmel P, Zhou H (2021) X-shaped structure of bacterial heterotetrameric tRNA synthetase suggests cryptic prokaryote functions and a rationale for synthetase classifications. Nucleic Acids Res 49:10106–10119.

28. Smith TF, Hartman H (2015) The evolution of Class II Aminoacyl-tRNA synthetases and the first code. FEBS Lett 589:3499–3507.

29. Valencia-Sánchez MI, Rodríguez-Hernández A, Ferreira R, Santamaría-Suárez HA, Arciniega M, Dock-Bregeon A-C, Moras D, Beinsteiner B, Mertens H, Svergun D, Brieba LG, Grøtli M, Torres-Larios A (2016) Structural Insights into the Polyphyletic Origins of Glycyl tRNA Synthetases. J Biol Chem 291:14430–14446.

30. Moody ERR, Mahendrarajah TA, Dombrowski N, Clark JW, Petitjean C, Offre P, Szöllősi GJ, Spang A, Williams TA (2022) An estimate of the deepest branches of the tree of life from ancient vertically evolving genes. eLife 11.

31. Achsel T, Stark H, Lührmann R (2001) The Sm domain is an ancient RNA-binding motif with oligo(U) specificity. Proc Natl Acad Sci USA 98:3685–3689.

32. Moody ER, Mahendrarajah TA, Dombrowski N, Clark JW, Petitjean C, Offre P, Szöllősi GJ, Spang A, Williams TA (2022) An estimate of the deepest branches of the tree of life from ancient vertically evolving genes. eLife 11:e66695.

33. Naganuma M, Sekine S, Fukunaga R, Yokoyama S (2009) Unique protein architecture of alanyl-tRNA synthetase for aminoacylation, editing, and dimerization. Proc Natl Acad Sci USA 106:8489–8494.

34. Yu Z, Wu Z, Li Y, Hao Q, Cao X, Blaha GM, Lin J, Lu G (2023) Structural basis of a two-step tRNA recognition mechanism for plastid glycyl-tRNA synthetase. Nucleic Acids Res 51:4000–4011.

35. Chandonia JM, Fox NK, Brenner SE (2017) SCOPe: Manual Curation and Artifact Removal in the Structural Classification of Proteins - extended Database. J Mol Biol 429:348–355.

36. Schaeffer RD, Liao Y, Cheng H, Grishin NV (2017) ECOD: new developments in the evolutionary classification of domains. Nucleic Acids Res 45:D296–d302.

37. Naganuma M, Sekine S, Chong YE, Guo M, Yang XL, Gamper H, Hou YM, Schimmel P, Yokoyama S (2014) The selective tRNA aminoacylation mechanism based on a single G•U pair. Nature 510:507–511.

38. Kumar A, Åqvist J, Satpati P (2019) Principles of tRNA(Ala) Selection by Alanyl-tRNA Synthetase Based on the Critical G3·U70 Base Pair. ACS Omega 4:15539–15548.

39. Penev PI, Alvarez-Carreño C, Smith E, Petrov AS, Williams LD (2021) TwinCons: Conservation score for uncovering deep sequence similarity and divergence. PLoS Comput Biol 17:e1009541.

40. McClain WH, Foss K, Jenkins R, Schneider J (1991) Four sites in the acceptor helix and one site in the variable pocket of tRNA (Ala) determine the molecule’s acceptor identity. Proc Natl Acad Sci USA 88:9272–9276.

41. Chong YE, Guo M, Yang X-L, Kuhle B, Naganuma M, Sekine S-i, Yokoyama S, Schimmel P (2018) Distinct ways of G: U recognition by conserved tRNA binding motifs. Proc Natl Acad Sci USA 115:7527–7532.

42. Nagato Y, Yamashita S, Ohashi A, Furukawa H, Takai K, Tomita K, Tomikawa C (2023) Mechanism of tRNA recognition by heterotetrameric glycyl-tRNA synthetase from lactic acid bacteria. The Journal of Biochemistry:mvad043.

43. Mohan S, Hsiao C, VanDeusen H, Gallagher R, Krohn E, Kalahar B, Wartell RM, Williams LD (2009) Mechanism of RNA double helix-propagation at atomic resolution. J Phys Chem B 113:2614–2623.

44. Vishwanath P, Favaretto P, Hartman H, Mohr SC, Smith TF (2004) Ribosomal protein-sequence block structure suggests complex prokaryotic evolution with implications for the origin of eukaryotes. Mol Phylogenet Evol 33:615–625.

45. Altschul SF, Gish W, Miller W, Myers EW, Lipman DJ (1990) Basic local alignment search tool. J Mol Biol 215:403–410.

46. Frickey T, Lupas A (2004) CLANS: a Java application for visualizing protein families based on pairwise similarity. Bioinformatics 20:3702–3704.

47. Steinegger M, Meier M, Mirdita M, Vöhringer H, Haunsberger SJ, Söding J (2019) HH-suite3 for fast remote homology detection and deep protein annotation. BMC Bioinformatics 20:473.

48. Swairjo MA, Otero FJ, Yang X-L, Lovato MA, Skene RJ, McRee DE, de Pouplana LR, Schimmel P (2004) Alanyl-tRNA synthetase crystal structure and design for acceptor-stem recognition. Mol Cell 13:829–841.

49. Antika TR, Chrestella DJ, Tseng YK, Yeh YH, Hsiao CD, Wang CC (2023) A naturally occurring mini-alanyl-tRNA synthetase. Commun Biol 6:314.

50. Frenkel-Pinter M, Petrov AS, Matange K, Travisano M, Glass JB, Williams LD (2022) Adaptation and Exaptation: From Small Molecules to Feathers. J Mol Evol 90:166–175.

51. Schimmel P, Yang XL (2004) Two classes give lessons about CCA. Nat Struct Mol Biol 11:807–808.

52. Tomita K, Fukai S, Ishitani R, Ueda T, Takeuchi N, Vassylyev DG, Nureki O (2004) Structural basis for template-independent RNA polymerization. Nature 430:700–704.

53. Westhof E, Dumas P, Moras D (1988) Restrained refinement of two crystalline forms of yeast aspartic acid and phenylalanine transfer RNA crystals. Acta Crystallogr A 44 (Pt 2):112–123.

54. Schimmel P, Giegé R, Moras D, Yokoyama S (1993) An operational RNA code for amino acids and possible relationship to genetic code. Proc Natl Acad Sci USA 90:8763–8768.

55. Schimmel P, de Pouplana LR (1995) Transfer RNA: from minihelix to genetic code. Cell 81:983–986.

56. Keng T, Webster TA, Sauer RT, Schimmel P (1982) Gene for Escherichia coli glycyl-tRNA synthetase has tandem subunit coding regions in the same reading frame. J Biol Chem 257:12503–12508.

57. Wagar EA, Giese MJ, Yasin B, Pang M (1995) The glycyl-tRNA synthetase of Chlamydia trachomatis. J Bacteriol 177:5179–5185.

58. Arutaki M, Kurihara R, Matsuoka T, Inami A, Tokunaga K, Ohno T, Takahashi H, Takano H, Ando T, Mutsuro-Aoki H (2020) G: U-independent RNA minihelix aminoacylation by Nanoarchaeum equitans alanyl-tRNA synthetase: an insight into the evolution of aminoacyl-tRNA synthetases. J Mol Evol 88:501–509.

59. Parker ET, Cleaves HJ, Dworkin JP, Glavin DP, Callahan M, Aubrey A, Lazcano A, Bada JL (2011) Primordial synthesis of amines and amino acids in a 1958 Miller H2S-rich spark discharge experiment. Proc Natl Acad Sci USA 108:5526–5531.

60. Botta O, Glavin DP, Kminek G, Bada JL (2002) Relative amino acid concentrations as a signature for parent body processes of carbonaceous chondrites. Orig Life Evol Biosph 32:143–163.

61. Lussi YC, Magrane M, Martin MJ, Orchard S, Consortium U (2023) Searching and Navigating UniProt Databases. Current Protocols 3:e700.

62. Capella-Gutiérrez S, Silla-Martínez JM, Gabaldón T (2009) trimAl: a tool for automated alignment trimming in large-scale phylogenetic analyses. Bioinformatics 25:1972–1973.

63. Guindon S, Dufayard JF, Lefort V, Anisimova M, Hordijk W, Gascuel O (2010) New algorithms and methods to estimate maximum-likelihood phylogenies: assessing the performance of PhyML 3.0. Syst Biol 59:307–321.

64. Lefort V, Longueville JE, Gascuel O (2017) SMS: Smart Model Selection in PhyML. Mol Biol Evol 34:2422–2424.

65. Anisimova M, Gil M, Dufayard JF, Dessimoz C, Gascuel O (2011) Survey of branch support methods demonstrates accuracy, power, and robustness of fast likelihood-based approximation schemes. Syst Biol 60:685–699.

