## Supplemental figures S1-S8 for "Common evolutionary origins of the bacterial glycyl tRNA synthetase and alanyl tRNA synthetase"

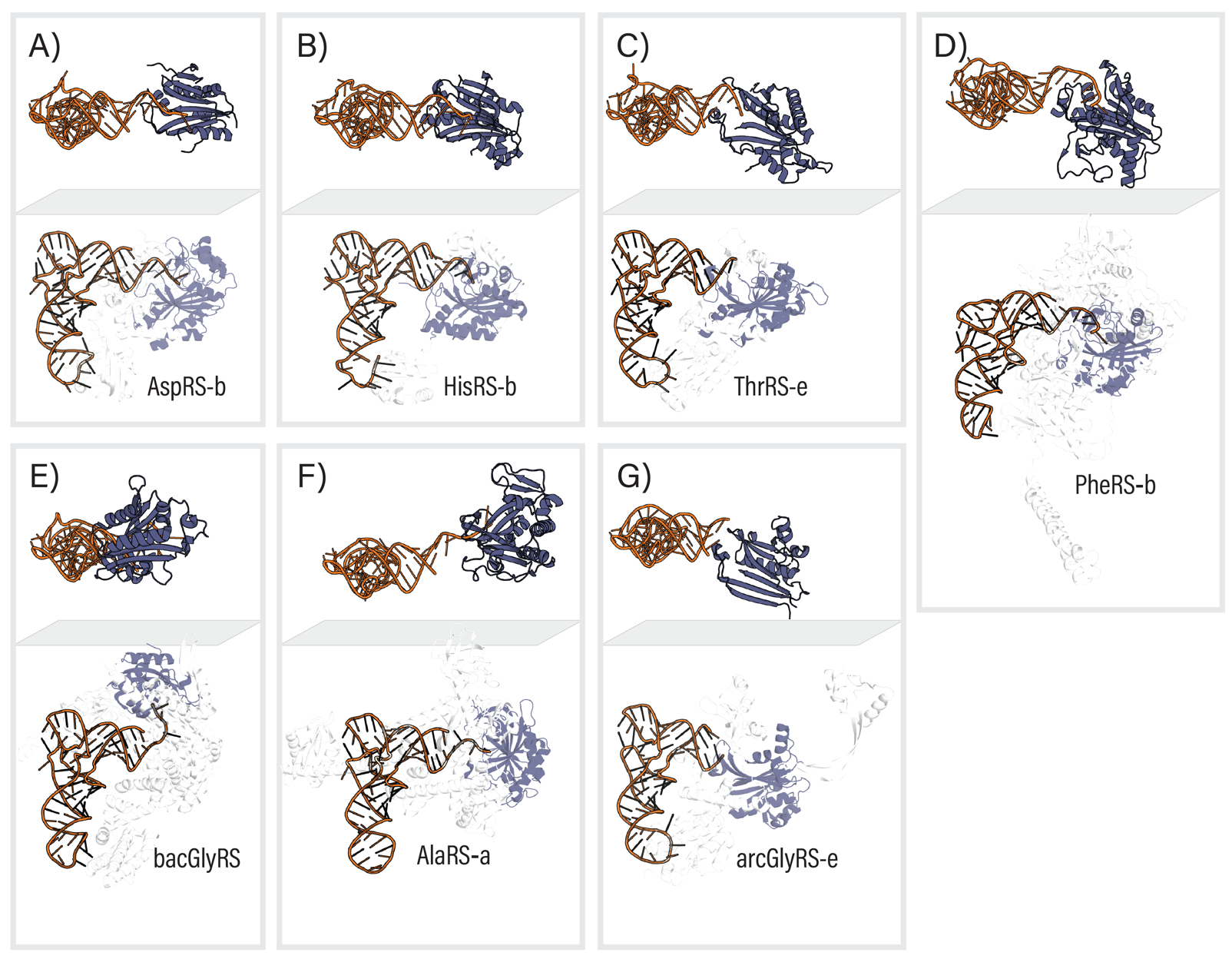

**Figure S1. Interaction between the catalytic domain and the CCA tail of tRNA in class II aminoacyl tRNA synthetases**. Top and lateral views of experimentally determined class II aminoacyl tRNA synthetases in complex with tRNA. Orange, tRNA molecule. Dark blue, catalytic domain. (A) AspRS from *Escherichia coli* (PDB 1C0A). (B) HisRS from *Thermus thermophilus* (PDB 4RDX). (C) ThrRS from *Saccharomyces cerevisiae* (PDB 4YYE). (D) PheRS from *Thermus thermophilus* (PDB 1EIY). (E) bacGlyRS from Escherichia coli (PDB 7YSE). (F) AlaRS from Archaeoglobus fulgidus (PDB 3WQY). (G) arcGlyRS from Homo sapiens (PDB 4QEI).

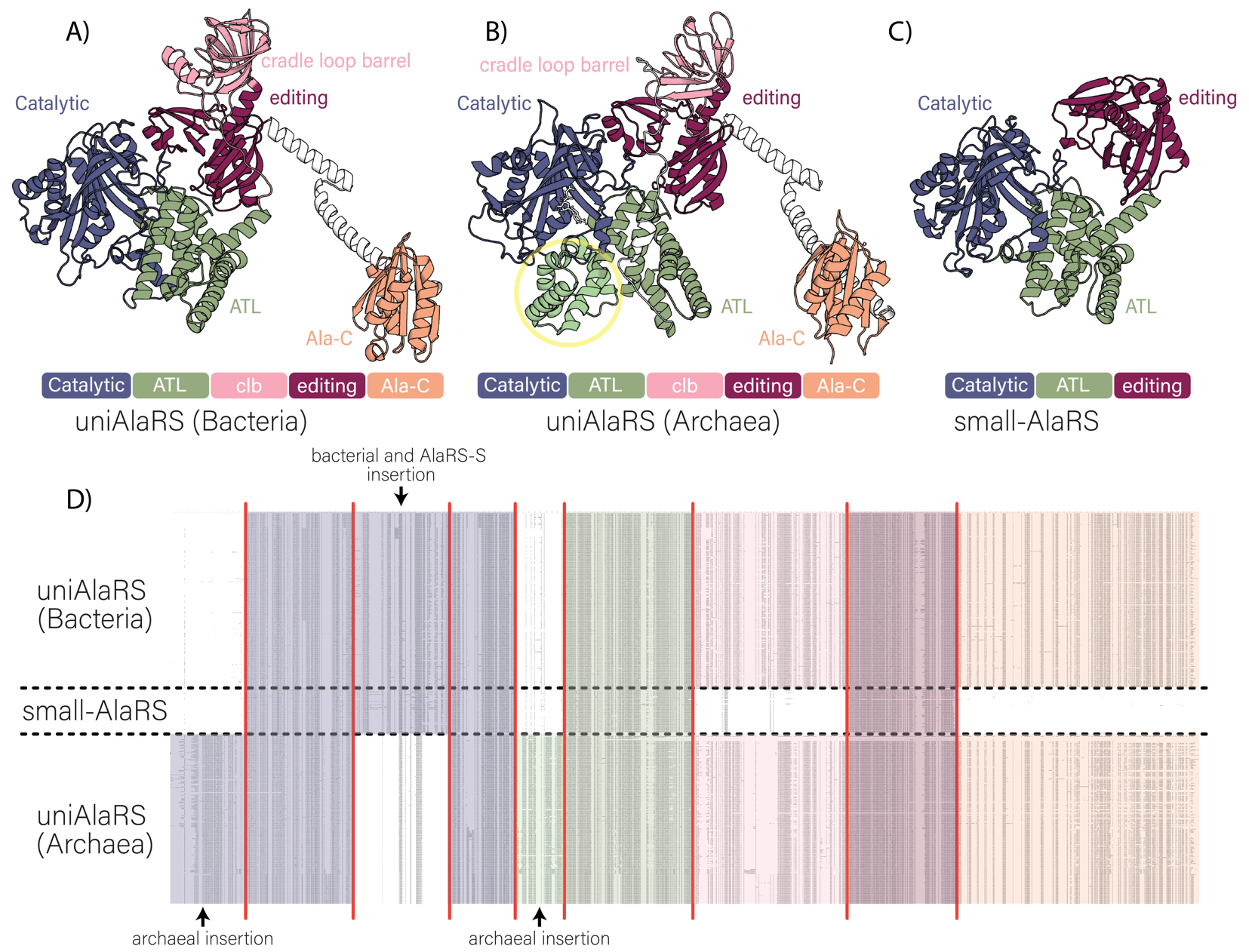

**Figure S2. Multi-domain organization of uniAlaRS and smallAlaRS.** (A) Structure and multidomain organization of uniAlaRS in *Aquifex aeolicus* (UniProt ID: O67323, AlphaFold ID: AF-O67323-F1). (B) Structure and multidomain organization of uniAlaRS in *Archaeoglobus fulgidus DSM 4304* (UniProt ID: O28029, PDB: 3WQY). (C) Structure and multidomain organization of small-AlaRS in *Thorarchaeota archaeon* (strain OWC) (UniProt ID: A0A524G8V1, AlphaFold ID: AF-A0A524G8V1-F1). (D) Multiple sequence alignment (MSA) shows the characteristic block structure between bacterial and archaeal sequences, as well as a third group of sequences that lack the cradle loop barrel and Ala-C domains.

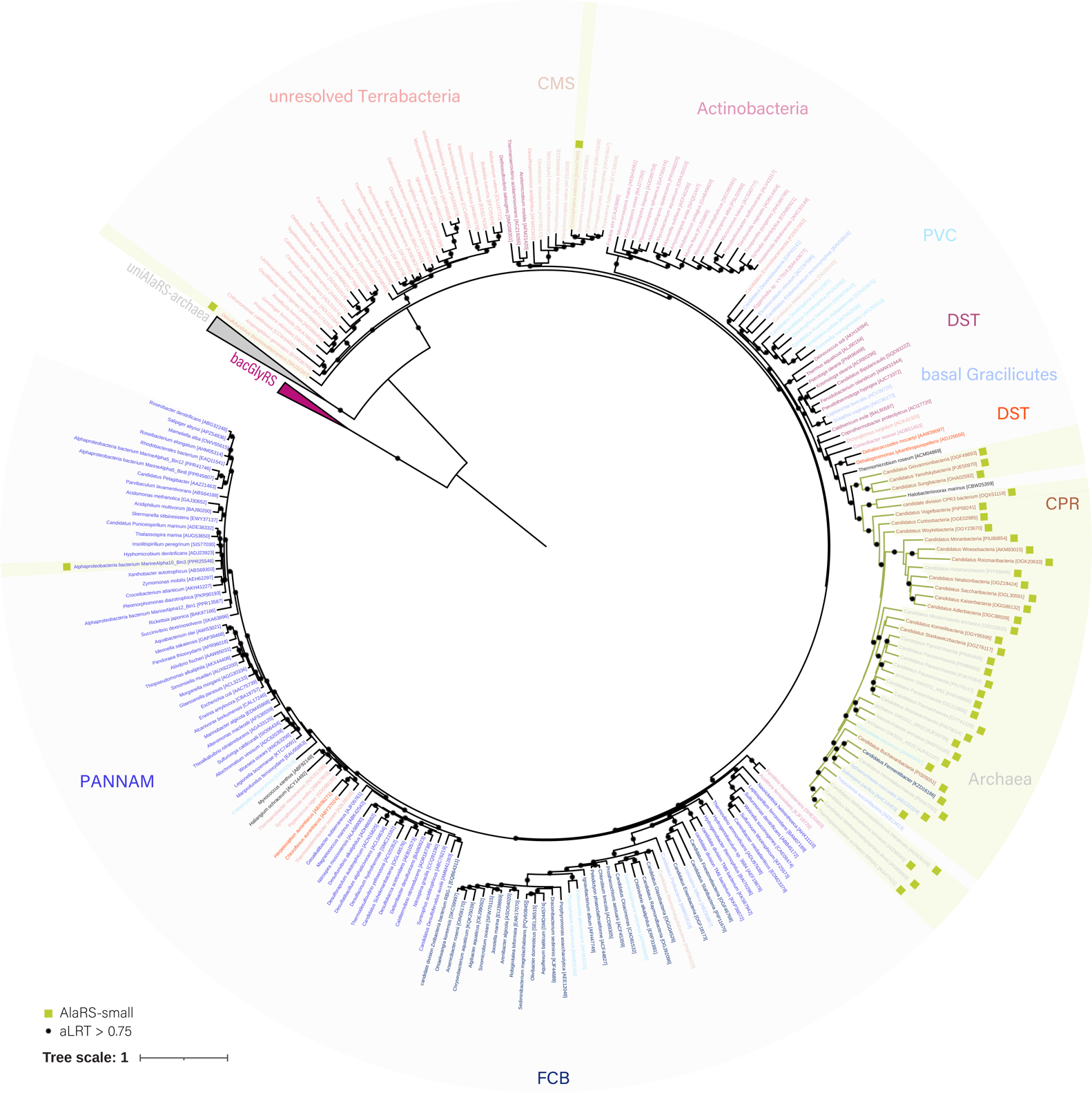

**Figure S3. Maximum-likelihood tree of catalytic and ATL domains including bacGlyRS and uniAlaRS sequences rooted at the midpoint.** The group of bacGlyRS homologs is collapsed. Most archaeal uniAlaRS form a single group. small-AlaRS homologs lack the Alanine Racemace C and C-Ala domains of canonical uniAlaRSs. Archaeal small-AlaRS sequences group with bacterial small-AlaRS sequences.

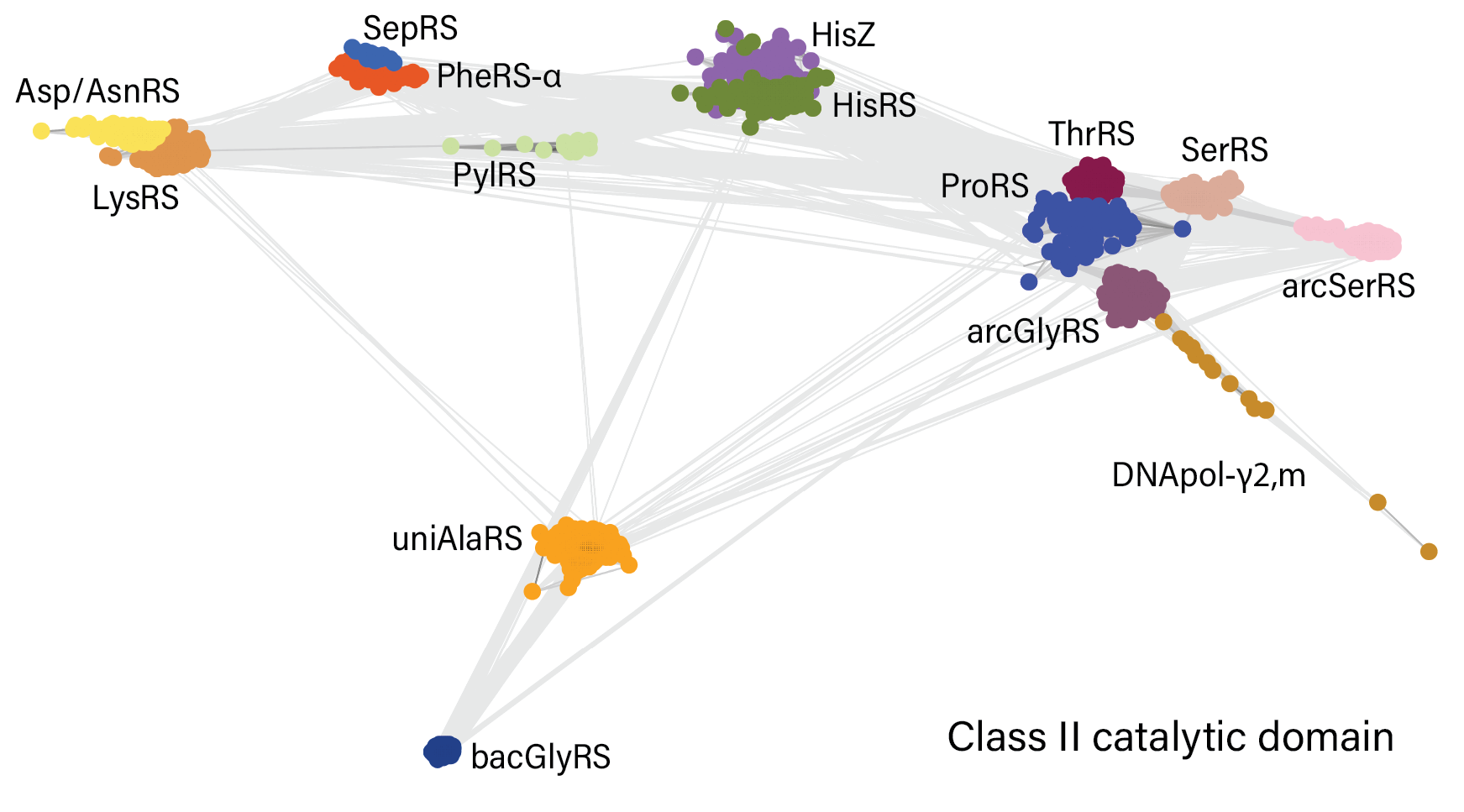

**Figure S4. Cluster map based on pairwise sequence similarities between class II catalytic domains.** Sequences of tRNA synthetases (Asp/AsnRS: Aspartate--tRNA(Asp/Asn) ligase; LysRS: Lysine--tRNA ligase; SepRS: O-phosphoserine--tRNA(Cys) ligase; PheRS-α: Phenylalanine--tRNA ligase subunit alpha; PylRS: Pyrrolysine--tRNA ligase; HisRS: Histidine--tRNA ligase; ProRS: Proline--tRNA ligase; ThrRS: Threonine--tRNA ligase; SerRS: Serine--tRNA ligase; arcSerRS: Type-2 serine--tRNA ligase; arcGlyRS: Glycine--tRNA ligase; and Alanine--tRNA ligase) and other proteins (HisZ: ATP phosphoribosyltransferase regulatory subunit and DNApol-γ2,m: DNA polymerase subunit gamma-2, mitochondrial) retrieved from the second sequence similarity search were trimmed to the class II catalytic domain and clustered with CLANS based on pairwise sequence similarities. Nodes represent class II catalytic domain sequences; connections between nodes indicate BLAST matches between pairs of sequences with a P-value lower than 1×10^-10^. Proteins of the same type are indicated by the same color. At a P-value of 1×10^-13^ the catalytic domain of bacGlyRS shows sequence relationships with only the catalytic domain of uniAlaRS.

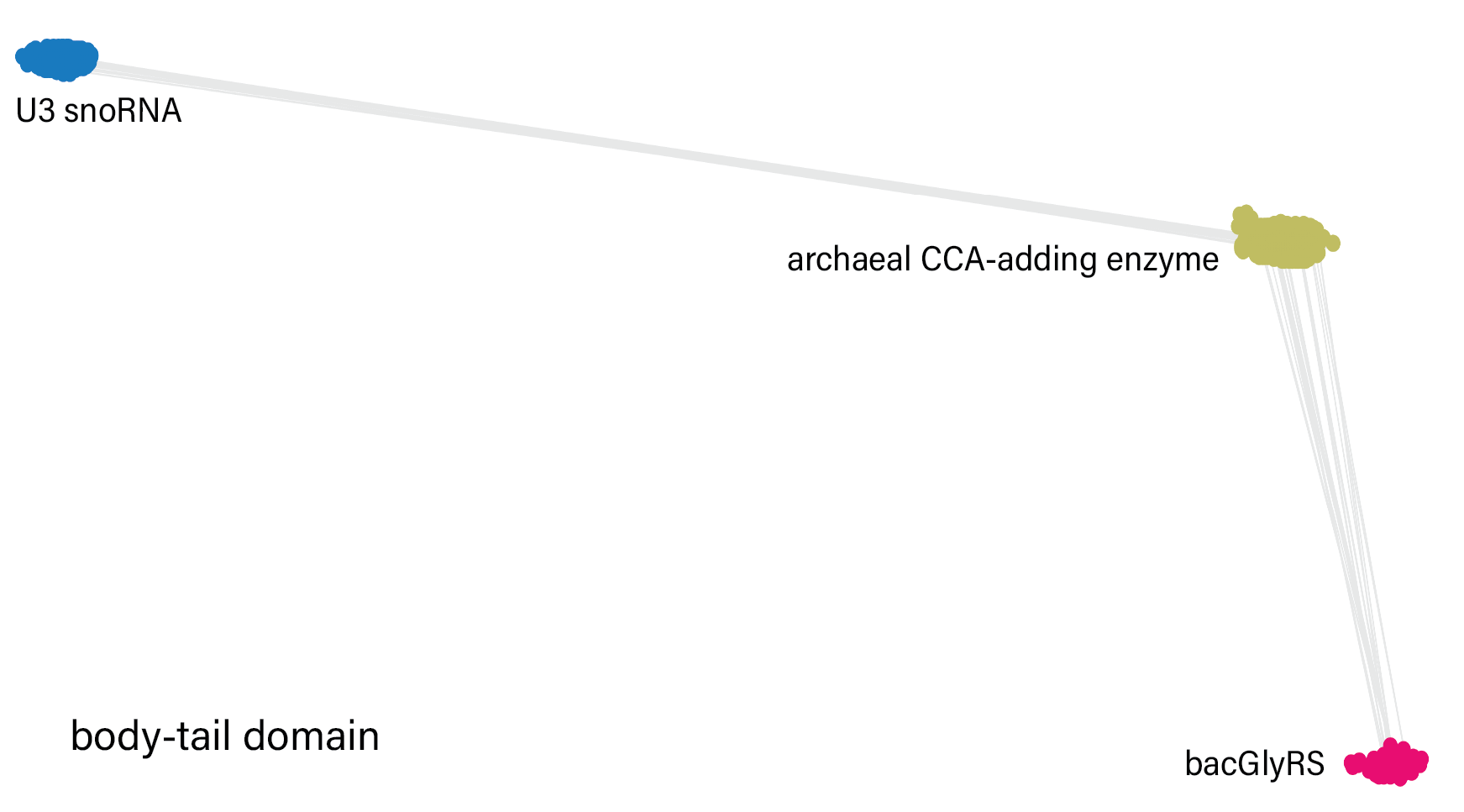

**Figure S5. Cluster map based on pairwise sequence similarities between body-tail domains.** Sequences of the eukaryotic U3 small nucleolar RNA-associated protein 22 (U3 snoRNA) and the archaeal CCA-adding enzyme retrieved from the second sequence similarity search were trimmed to the body and tail domains and clustered with CLANS based on pairwise sequence similarities. Nodes represent body-tail domain sequences; connections between nodes indicate BLAST matches between pairs of sequences with a P-value lower than 1×10^-10^. Proteins of the same type are indicated by the same color. At a P-value of 1×10^-10^ the body and tail domains of bacGlyRS shows sequence relationships with only the body and tail domains of the archaeal CCA-adding enzyme.

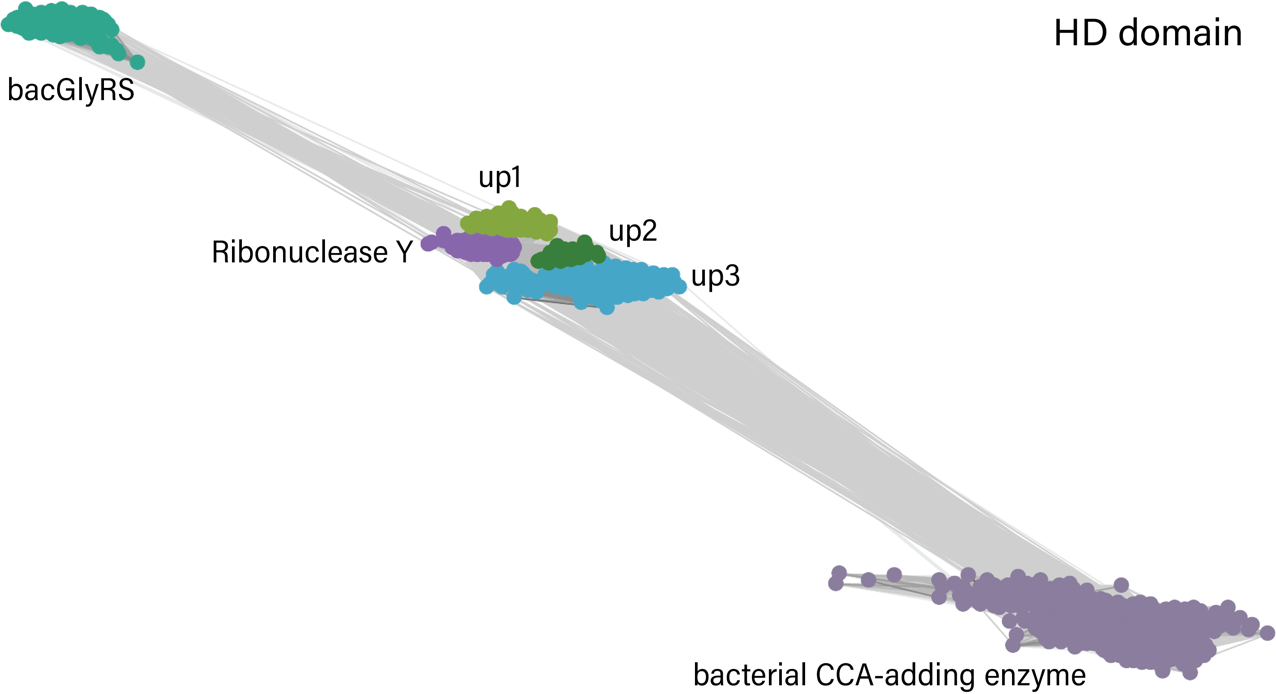

**Figure S6. Cluster map based on pairwise sequence similarities between HD domains.** Sequences of Ribonuclease Y, the bacteria CCA-adding enzyme and several uncharacterized proteins (up1; up2; and up3) retrieved from the second sequence similarity search were trimmed to the body and tail domains and clustered with CLANS based on pairwise sequence similarities. Nodes represent HD domain sequences; connections between nodes indicate BLAST matches between pairs of sequences with a P-value lower than 1×10^-10^. Proteins of the same type are indicated by the same color. At a P-value of 1×10^-13^ the HD domain of bacGlyRS sequences show relationships with only the HD domain of Ribonuclease Y (GenBank identifiers: SME90275.1; PKM99705.1; PIV64412.1; OSM07205.1; BAJ31136.1; SMG21051.1; OGM04846.1; OGI11153.1; and EDM29070.1) and with uncharacterized proteins ABY35864.1 and ABQ90411.1.

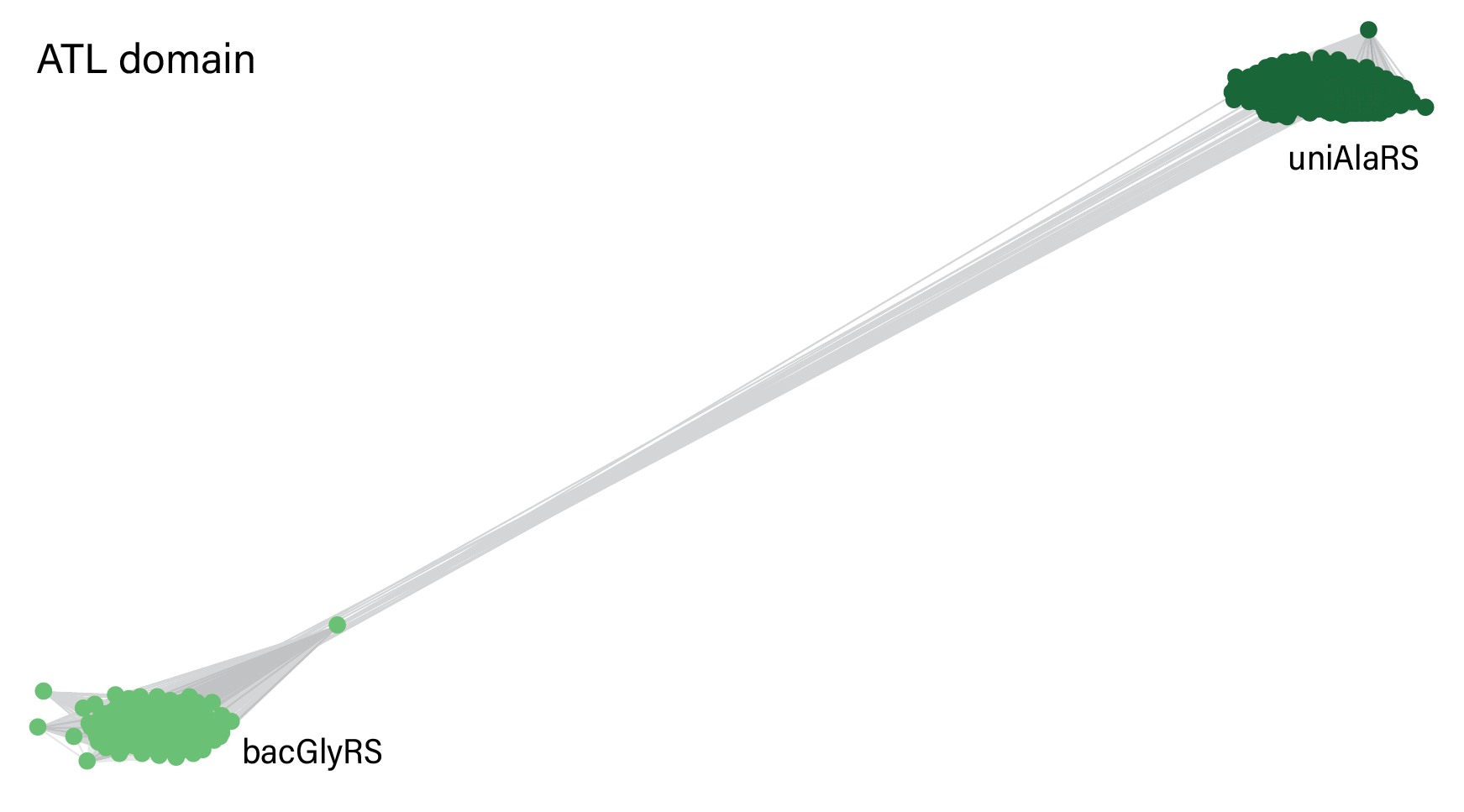

**Figure S7. Cluster map based on pairwise sequence similarities between ATL domains.** Sequences of uniAlaRS retrieved from the second sequence similarity search were trimmed to the ATL domain and clustered with CLANS based on pairwise sequence similarities. Nodes represent ATL domain sequences; connections between nodes indicate BLAST matches between pairs of sequences with a P-value lower than 1×10^-10^. Proteins of the same type are indicated by the same color.

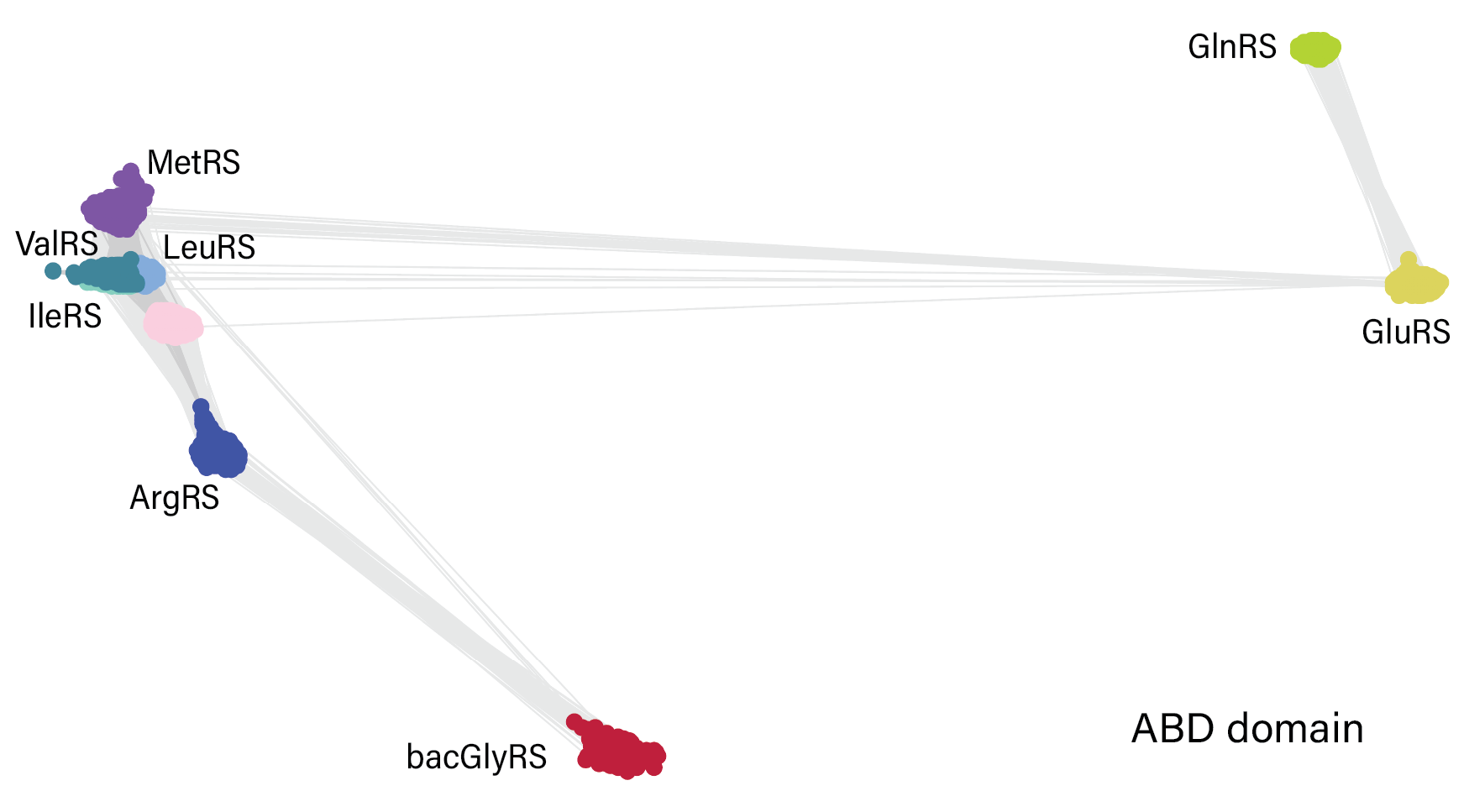

**Figure S8. Cluster map based on pairwise sequence similarities between ABD domains.** Sequences of tRNA synthetases (ArgRS: Arginine--tRNA ligase; IleRS: Isoleucine--tRNA ligase; VelRS: Valine--tRNA ligase; LeuRS: Leucine--tRNA ligase; MetRS: Methionine--tRNA ligase; GluRS: Glutamate--tRNA ligase; and GlnRS: Glutamine--tRNA ligase). Nodes represent ABD domain sequences; connections between nodes indicate BLAST matches between pairs of sequences with a P-value lower than 1×10^-10^. Proteins of the same type are indicated by the same color. At a P-value of 1×10^-12^ the body and tail domains of bacGlyRS shows sequence relationships with only the ABD of ArgRS.
